# Gate keeping the sebaceous gland

**DOI:** 10.1101/2023.08.22.554243

**Authors:** Suveer Sachdeva, Araz Ahmed, Gordon Proctor, Isabelle Miletich

## Abstract

Sebaceous gland (SG) secretions are pivotal to skin and eye health. At the SG-epidermis confluence is an overlooked epidermal collar we named the follicular epidermis (FE). Here, we show that the FE is similar among different SG types and contains unique Axin2+ stem cells in a niche at its basal FE-perpendicular flexure (FE-PF). Lineage tracing and ablation assays demonstrate Axin2+ cells as integral for FE homeostasis. Wnt-secretion arrest experiments resulted in FE-obstruction via hyperproliferation and inhibited differentiation inferring FE-PF Axin2+ cells are Wnt-producers maintaining FE-patency. Upon constitutive Axin2+ cell Wnt signalling, FE-PF-specific cell proliferation with early-stage signs of malignancy formed alongside a keratin-plugging FE-obstruction type. While keratin-based obstructions are recognized, inhibited differentiation obstructions are not, which suggests a one-size-fits-all therapeutic approach is not optimal. We offer a molecular identification toolkit to aid anti-obstruction advances in dermatology/ophthalmology and highlight the FE-PF as a skin tumorigenic site.

**Summary:** This work characterises a novel site in the skin, it’s stem cells and niche, a site prone to cancer. Obstructions can occur at this site by two mechanisms, one of which is novel, bringing into question current therapy and prompting a rethink of disease in dermatology and ophthalmology to account for this novel site.

**Graphical Abstract:** 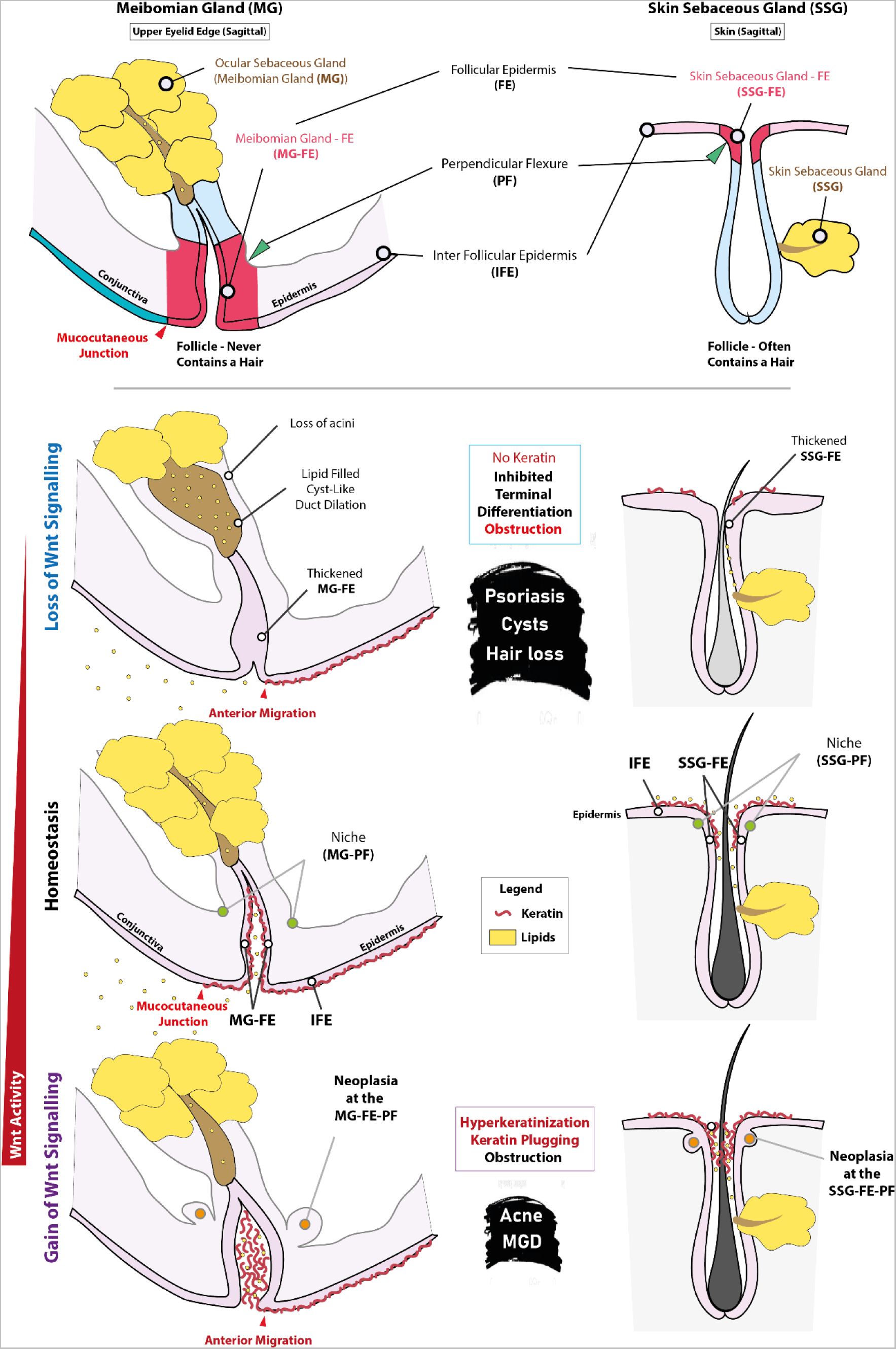

## Introduction

The skin is the largest organ of the body and critical for survival. The outer most layer is the interfollicular epidermis (IFE) and is tasked with barrier protection. The cornea performs barrier protection for the ocular globes, but is lined by corneal epithelium. Optimal function of both epithelia predicates on healthy sebaceous gland (SG) secretions. In the skin, the smaller skin SG (SSG) produces sebum, while in the eyelids, the largest SG, the meibomian gland (MG) produces meibum, and both are critical to body surface homeostasis ^1,2^.

The sebaceous follicle (SF), a small pouch-like secretory cavity, contains the SG. The SSG-SF is often associated with a hair, while the MG-SF never contains a hair. At the terminal end of both SFs lies a perpendicular invaginated collar of cylindrically shaped epidermis connecting the SF to the IFE. This site has not been attributed a name separate to the IFE although it is morphologically and functionally distinct, and vulnerable to obstructions that inhibit sebaceous secretions. It can be argued some recognition of this site has been made. In MG research, this site has been referred to as ‘the ampulla’ ^3^. In the skin hair-follicles it has been referred to as the ‘the oracro-infundibulum’ ^4^. Neither naming the epidermal collar itself. Recent notable work by Roy *et al.* (2016) showed that epidermal progenitor cells closer to a hair, in the dorsal skin of mice, experienced increased proliferation compared to those more distant. This is indicative of a difference in stem cell properties and their associated niches in the IFE. Herein, we refer to the collar of epidermis belonging to the follicle as the ‘follicular epidermis’ (FE).

Epidermal renewal occurs via basal stem/progenitor cell proliferation and differentiation to balance routine shedding of the keratinized cornified layer. As a cell differentiates, it moves apically entering the spinous layer. Here, the cell begins production of keratohyalin granules (keratin precursors). The cell then enters the granular layer where active keratin production begins. Upon terminal maturation, the cell nucleus and organelles are lost while keratin is accumulated and then shed ^5,6^. Keratin production adds stiffness, rigidity, smoothness, and resilience in skin ^7–9^. Keratin disorders and their pathophysiology are well documented ^10,11^ but lacking impact consideration at the FE.

Clinical pathological obstructions at the region of the meibomian gland-FE (MG-FE) and skin sebaceous gland-FE (SSG-FE) are commonplace with striking similarities, examples include meibomian gland dysfunction (MGD) and Acne vulgaris. MGD is commonly characterised by MG terminal-duct obstruction ^12^ due to hyperkeratinisation ^13–17^, with or without inflammation ^18–20^ and is suggested to affect 35.8% globally ^12,21^. Acne vulgaris is one of the most common skin diseases affecting approximately 85% of adolescents ^22,23^. Obstructions in acne involve plugging at the FE region, and current research emphasises the role of altered sebum characteristics, hormonal imbalances, bacterial influence, and inflammation in plugging ^24,25^, although a hyperkeratinisation pathophysiological component of the disease has previously been stated ^23,26^. Acne is known to cause permanent physical scarring, reduction in quality of life and is associated with increased rates of depression, anxiety, and suicidal ideation ^25^. Notably, drug-induced acne (DIA) is a common clinical condition linked to systemic drugs such as corticosteroids, lithium, vitamin B12, among several others ^27^. In both MGD and acne, the hyperkeratinisation involved remains poorly understood but is the only proposed epidermal obstruction mechanism. The causes of obstructions of both these diseases remain elusive, they are still better pathologically characterised than other diseases in their respective regions. Epidermal/sebaceous cysts are common conditions with an unknown cause. In the eyelid, MG cysts, also known as chalazions, are the most common inflammatory lesion of the eyelid ^28,29^, yet pathophysiology of these lesions remains vague with reports suggesting an obstructive component attributed to meibum alterations causing plugging ^29,30^. In the skin, epidermoid cysts are the most common cutaneous cyst. Epidermoid cysts are suggested to arise from ‘trapped keratin’ ^31^ but a mechanism as to how this occurs is lacking. Clinically, several other idiopathic conditions with obstructive links are known to occur in the skin, termed follicular occlusion disorders. These include but are not limited to hidradenitis suppurativa, folliculitis, keratosis pilaris, acne conglobata, acne rosacea, and dissecting cellulitis to name a few ^32–38^.

In homeostasis, the Wnt pathway’s secreted Wnt proteins are known to stimulate and regulate epidermal stem cell self-renewal, proliferation, differentiation, polarity, and migration ^39–41^. Wnt signalling is a major cue directing skin maintenance ^39,40^ and aberrant Wnt signalling has been linked to several diseases and cancer ^42–46^.

In light of all of the above, we hypothesised that Wnt signalling is influential in homeostasis of the FE-maintenance of SG patency and preventing obstructions. We used several transgenic mouse lines to interrogate Wnt signalling’s role in the FE of the advantageously large hairless MG-FE, and in the smaller (haired) eyelid SSG-FE, comparatively, to altogether assess SG FE apparati.

We show that stem/progenitors with Wnt activity reside in the FE basal layer at the perpendicular flexure (PF) in both SGs, a site that serves as a niche and a location where aggressive neoplasia specifically develops upon constitutive Wnt activation in FE stem/progenitor cells. Furthermore, we show that opposing Wnt activity manipulations results in mechanistically distinct obstructions, one of which mimics known keratinised obstructions, while the other lacks keratinisation and is currently unknown to occur. This latter finding offers valuable insight to current and potentially overlooked SG dysfunctions and their therapy. Altogether, our findings offer novel insight and potential therapeutic avenues for several frequent diseases in ophthalmology and dermatology, and, furthermore, we outline a region prone to skin cancer upon Wnt signalling deregulation, which may aid early diagnostics in oncology.

## Results

### 1. Global *Wls*-depletion results in FE-obstruction due to hyperplasia prior to inflammation

Recently Augustin et al. (2013) characterised mice with an epidermal conditional knock-out of the Wnt cargo receptor/protein *Wls* (Wntless/*Evi/Gpr177*). They found K14-cre *Wls*-depletion to result in hair loss, skin inflammation and impaired epidermal barrier function. *Wls*-down-regulation identified in human psoriatic skin biopsies led them to suggest that *Wls*-depletion mice developed epidermal lesions resembling human psoriasis.

The FE is currently not formally recognised but found terminally at the MG (MG-FE) and SSG (SSG-FE) **(Fig. 1Q)**. To determine the effects of *Wls*-depletion at the FE we used *pCagCreERT2/+; Wlsfl/fl* mice. Two weeks following global-*Wls*-depletion, gross phenotypes included visible eyelid inflammation and corneal opacification **(Fig. 1B, C)**. Oil-red-O wholemount staining revealed MG obstructions, central ductal dilation, acinar and eyelash loss **(Fig. 1D, E)**. Sagittal haematoxylin and eosin staining (H&E) revealed a hyperplastic obstruction in the MG-FE at 1-week with an apparent alteration in IFE cornification including antero-placement of the mucocutaneous junction (zone of keratinization – a clinical diagnostic of MGD as per the line of Marx) ^47–49^, acini loss, and central duct dilation and thickening **(Fig. 1F, G)**. 2-month samples showed MG-FE degeneration, cyst-like ductal dilation, likely linked to marked acinar loss, and a residual sign of clinical MGD **(Fig. 1H)**. Transverse sectioning and H&E allowed alternate visualization of the FE obstructions in both SGs **(Supp. 1A-X)**.

**Figure 1:**
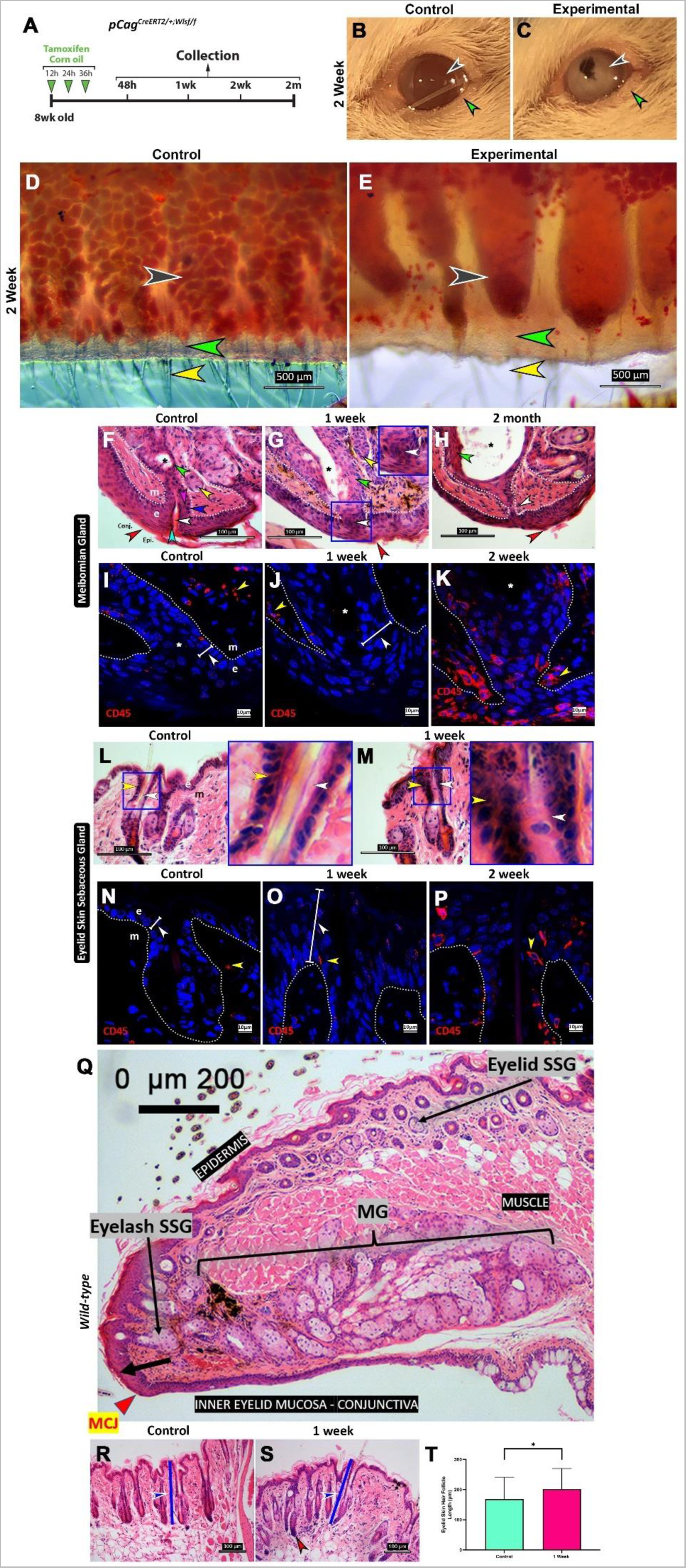
Global Wls-depletion results in FE-obstruction due to hyperplasia prior to inflammation. **A** Tamoxifen (experimental) and corn-oil (control) time course administration to 8-week-old *pCag^CreERT2/+^Wls^f/f^* littermate mice (n≥3 for all timepoints). **B** 2-week **c**ontrol mice have translucent corneas **(black)** with smooth, pink-coloured eyelids, eyelashes visible **(green)**. **C** Experimental have possible corneal opacification **(black). R**aised and pale eyelid appearance suggests inflammation with a loss of eyelashes **(green)**. **D** 2-week control whole mount Oil-red-O (ORO) of eyelids viewed from the inner aspect show lipids (red stain) within MG acini **(black),** healthy **MG-FE (green),** eyelashes **(yellow)**. **E** Experimental show loss of MG acini, lipid-filled central duct dilation **(black)**, abnormal MG-FE **(green)**, a loss of eyelashes **(yellow)**. **F-H, L,M** H&E on sagittal sections. F Control sample at MG-FE. Conjunctiva -**Conj.,** Epidermis – **Epi.,** red arrow indicates the keratinization terminating at the MCJ (pink crystal-like appearance of stratum corneum) is posterior to the MG-FE. Basal cells at the MG-FE have large, oval, nuclei with dense stain **(blue)**, above, smaller, lighter stained nuclei of spinous cells **(magenta)**, above, flattened, and gritty most opaque nuclei of the granular layer **(cyan),** and above, the cornified layer devoid of nuclei and filled with crystal like pink keratin **(white)**. Proximally a healthy MG central duct lumen **(asterisk)** and its epithelium (**green)**. Acinus present (**yellow)**. **G** 1-week experimental sample shows an MG-FE obstruction linked to hyperplasia with involved cell nuclei carrying a dense stain **(white)**. Proximally, central duct epithelial thickening **(green)**, luminal dilation **(asterisk)** and a loss of acini **(yellow)**. Blue square is MG-FE magnified inset. Zone of keratinisation is displaced anterior to the MG-FE **(red)** with overall appearance of reduced IFE cornification. **H** 2-month samples show a severely obstructed MG-FE **(white)**, central duct thickening present **(green)** with increased ductal dilation **(asterisk)**. Zone of keratinisation remains anteriorly displaced **(red)**. **I-K, N-P** Immunofluorescence for CD45+ immune cells at the respective FE. **I** Few immune cells seen in the MG-FE control and **(J)** at 1-week, although MG-FE obstruction is present **(white)**. **K** 2-week sample shows significant inflammation at the MG-FE. **L** 2-week control SSG-FE has thin epithelium **(yellow)** encapsulating a hair-shaft with space between the two **(white)**. M 1-week experimental samples show a thickened SSG-FE **(yellow)** with a loss of space between the epithelium and the hair shaft **(white)** i.e., obstruction. **N** Few immune cells seen in the SSG-FE control and **(O)** at 1-week, although SSG-FE thickening is present **(white)**. **P** 2-week sample shows significant inflammation at the SSG-FE. **Q** Sagittal H&E of an upper wild-type mouse eyelid showing the location and relative size of the MG and SSG. **R** Control shows hair-follicles of normal length in the dermal portion (blue) **S** 1-week samples show elongation of the dermal portion of the hair-follicle, particularly at the bulb region (red). **T** Quantification of the increase in length of the dermal portion of the hair-follicle. Scale bars per image are indicated.

Inflammatory cell involvement was then assessed via immunofluorescence for CD45, the pan-immune cell marker. While the hyperplastic obstruction was well established at 1-week, immune cell numbers only appeared significant at the 2-week timepoint suggesting inflammation to be secondary to the obstruction **(Fig. 1I-K)**. In the SSG-FE at 1-week, a loss of space was detected between hair-shafts and their adjacent epithelia, which had marked thickening **(Fig. 1L, M).** CD45+ inflammatory cell detection resembled findings in the MG-FE **(Fig. 1N, P)**. At 1 week, the hairs within the SSG follicle were found to have an increased dermal length particularly at the bulb region of the hair **(Fig. 1R-T)**. Altogether it suggests that *Wls*-depletion results in MG and SSG obstructions associated with hyperplasia and secondary inflammation, and the hair-follicles of the SSG had undergone dermal lengthening.

### 2. SG obstruction following global *Wls*-depletion features inhibited differentiation

We then interrogated the mechanism of the obstruction present upon *Wls*-depletion using key molecular markers. First, MG-FE proliferation was assessed using the nuclear marker of proliferation Ki67 with cytokeratin-5 (K5), a basal cell marker. *Wls*-depletion caused an increase in basal Ki67+ proliferating cell numbers from 7±2% to 37±3% at 48-hours and from 7±1% to 55±4% at 1-week. The excess proliferation state subsided at 2-months with control at 8±1% and experimental at 10±2% **(Fig. 2 A-D, Q)**.

**Figure 2.**
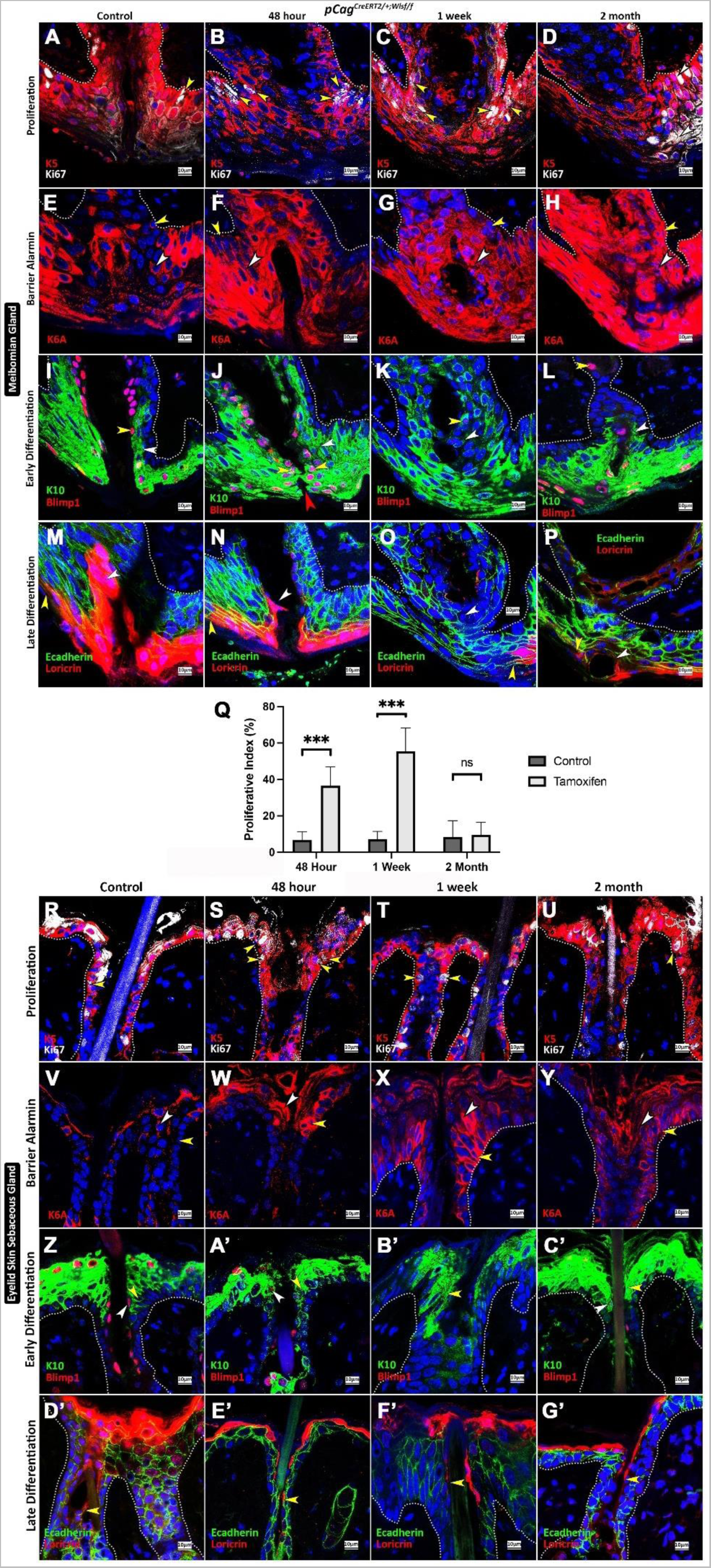
Global Wls-depletion FE-obstruction features inhibited differentiation. 8-week-old *pCag^CreERT2/+^Wls^f/f^*littermate mice injected with tamoxifen (experimental) or corn oil (control) (n≥3 for all timepoints). Immunofluorescence at the **A-Q** MG-FE and **R-G’ SSG-FE**. **A** Shows healthy Ki67+ proliferating cells **(yellow)** in the basal, K5+ MG-FE. An increase in proliferation in 48-hour **(B)** and 1-week samples. **D** Shows a decrease in proliferation in the 2-month sample. Quantification of proliferation shown in **Q**. All counts were derived from n=3 animals per timepoint and subjected to unpaired student t-tests (error bars = SD). **E** K6A expression is sparsely present in the basal **(yellow)** and suprabasal **(white)** layer of the MG-FE. **F-H** 48-hour to 2-month samples showed a progressive increase in K6A expression. **I** K10+Blimp1-expression labels the spinous and K10+Blimp1+ labels granular layers **(white)** in homeostasis. Blimp1+ labels terminally differentiated cell nuclei of the granular layer **(yellow)**. **J** 48-hour shows an increase in K10 and Blimp1 cell numbers with opposing MG-FE Blimp1+ cells converging to obstruct **(red)**. **K** 1-week shows marked reduction in Blimp1 expression **(yellow)** and presence of K10+ cells luminally **(white)**. **L** 2-month shows, a MG-FE with reduction of K10 and Blimp1 expression. **M** Loricrin labels the granular and cornified layers of the MG-FE extending proximally into the lumen **(white)** also indicating the MCJ keratinization **(yellow)** posterior to the MG-FE in homeostasis. **N** 48-hour Loricrin expression was reduced within the MG-FE lumen **(white arrow)**, somewhat normal zone of keratinisation **(yellow arrow)**. **O** 1-week shows marked loss of Loricrin expression with anterior displacement - clinical sign of MGD. **P** 2-month shows continued defective Loricrin expression. Scale bars and markers per image are indicated. **R-G’** Shows similar if not identical findings in the SSG-FE.

Recent work suggests Keratin-6 (K6) serves as a crucial early barrier alarmin whose upregulation results in hyperproliferation while altering keratinocyte inflammatory features ^50–52^. We found *Wls*-depletion resulted in an increase in K6A+ expression at the MG-FE **(Fig. 2E-H)**. We then investigated differentiated cell layers where K10+Blimp1-labels cells in the early differentiated spinous layer, while Blimp1 labels nuclei of cells in the granular layer. At 48-hours we found an increase in spinous cell numbers with the granular layer of opposing margins of the MG-FE leading the obstruction. 1-week spinous cell numbers remained high with a loss of the granular layer thus linking spinous cells to the hyperplasia. At 2-months a marked loss of expression was seen among a diseased MG-FE **(Fig. 2I-L).** We then assessed the cornified layer using Loricrin, a structural protein present in terminally differentiated cells. Loricrin comprises more than 70% of the cornified envelope ^53^ allowing visualization of the MCJ and the terminal lumen of the MG-FE. A progressive reduction in Loricrin expression was seen at the MG-FE with permanent clinical signs of MGD setting in at 1-week **(Fig. 2M-P)**. Similar observations were made in the SSG-FE **(Fig. 2R-G’)**.

Altogether, *Wls* depletion appears to cause FE-obstruction by hyperproliferation, pathogenic K6A expression, hyperplasia in the spinous layer, and a loss of terminal differentiation i.e., cornification commencing at the granular layer.

### 3. A single stem/progenitor cell population with Wnt activity resides in a niche at the FE-PF

Given that the FE is morphologically, functionally, and molecularly distinct to the IFE, and that onset of an FE-obstruction upon *Wls*-depletion is rapid, we hypothesised that the FE has independent renewal characteristics involving Wnt-active stem/progenitor cells. To search for a likely niche-location, we first considered histology of pig and wild-type mouse FEs for clues. A raised-pouch-like basal region is always seen in pig samples (n>3) at the point of the basal FE-PF in both the MG and SSG **(Fig. 3A, D),** while sometimes less discernible in the mouse **(Fig. 3B, E, U)**. The PF is close by, basal, protected and at a point of flexure where it is likely exposed to unique mechanical cues. When employing the *TCF/Lef:H2B-GFP* mouse line ^54^, a faithful single-cell indicator of active Wnt signalling, rare cells were detected specifically at the PF **(Fig. 3C, F)**, earmarking this region as a likely-niche.

**Figure 3.**
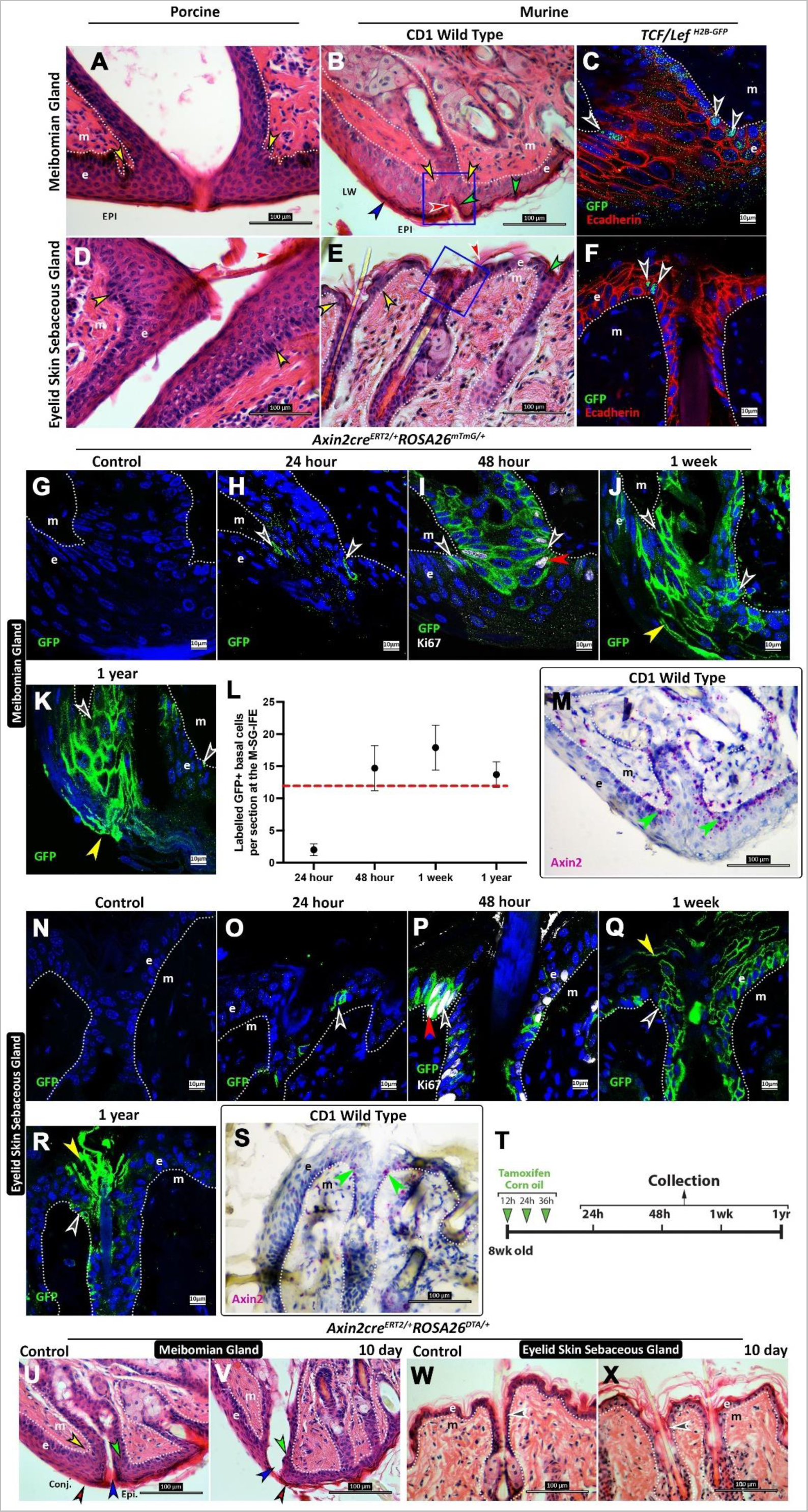
A single Wnt stem/progenitor cell population resides in a niche at the FE-PF. Sagittal H&E of pig **A** MG-FE and **D** SSG-FE show an irregular/raised basal layer at the FE-PF **(yellow).** Mouse **B** MG-FE and **E** SSG-FE are similar. *TCF/Lef:H2B-GFP* mice show Wnt active cells to specifically reside at the **C** MG-PF **F** SSG-PF **(black). T** Tamoxifen (experimental) and corn-oil (control) time course administration to 8-week-old Axin2^creERT2+^;ROSA26^mTmG/+^ littermate mice (n≥3 for all timepoints) to enable **G-K,** MG, **and N-R** immuno-fluorescent lineage tracing. **G, N** No control line leakage. **H, O** 24-hour Axin2 clones seen at the SSG-PF **(black)**. **I, P** Clonal expansion at 48-hours some GFP+ cells proliferating (Ki67+) **(red)**. **J, Q** 1-week Axin2+ progeny seen renewing whole SSG-FE epithelium **(yellow). K, R** 1-year Axin2 cells are long-lived continually serving renewal. **L** Quantification of MG-FE Axin2 basal clones (error bars = SD, n≥3 mice). **M, S** RNA scope for Axin2 on WT mice shows expression restricted basally at the SSG-FE **(green)**. **U-X** Sagittal H&E of following Axin2 ablation via *Axin2^CreERT2/+^ROSA26^DTA/+^* mice (n=3) with littermate corn oil controls, collected at day 10. **U** Control conjunctiva **(conj.)** meets the epidermis **(epi.)** where cornification terminates posterior to the MG-FE **(red)**, healthy cornification **(blue)**, and epithelial thickness **(green)** within MG-FE. Raised pouch-like appearance at MG-PF in this control sample **(yellow)**. **V** 10-day shows anterior displacement of cornification **(red and blue)** and a loss of epithelial cell layers within the MG-FE **(green). W** Control SSG-FE shows healthy epithelial thickness with containing dense nuclei **(black)**. **X** 10-day shows loss of epithelial thickness and fewer nuclei **(black)**. All counts were derived from n=3 animals per timepoint and subjected to unpaired student t-tests. Scale bars and markers per image are indicated.

Recently, Lim et al. (2013) studied the mouse plantar epidermis, a site devoid of SGs. They suggested a single stem/progenitor cell population that generates their own self-renewing autocrine Wnt signals to be responsible for plantar-IFE maintenance. Furthermore, they suggested long-range paracrine Wnt inhibitor secretion to guide progeny differentiation (Lim et al., 2013). These findings highlight a critical role for Wnt signalling in plantar-IFE homeostasis. Thus, we then performed a lineage tracing assay using *Axin2^creERT2+^;ROSA26^mTmG/+^*mice to identify Axin2+ Wnt active cells and their clonal progeny over key timepoints. At 24-hours, Axin2+ cells were seen at the FE-PF. At 48-hours, progeny expansion had occurred, at 1-week renewing the whole FE epithelium. Renewal was seen up to 1-year indicating the Axin2+ cells are long-lived and have key stem/progenitor characteristics of the FE in both SGs **(Fig. 3H-K, O-R)**. Number of basal clones were counted at each time point in the MG-FE and appeared to maintain stable numbers over time **(Fig. 3L)**. Approximately 37±1% of the Axin2+ clones were found to be proliferating at 48 hours **(S2 I)**. RNAscope for Axin2 on wild-type mice confirmed the PFs of both SGs as a region of marked expression **(Fig. 3M, S)**.

To assess the importance of Axin2+ clones, Axin2+ cell ablation using *Axin2^CreERT2/+^ROSA26^DTA/+^* mice was performed. At day 10, ablation resulted in a characteristic loss of FE morphology **(Fig. 3 U-X) (S2 A-F, G, H, J, K)** and a marked reduction in proliferation **(S2 L-P)** in both SGs attesting to the importance of these cells.

### 4. Depleting *Wls* in Axin2+ stem/progenitors reveals them as critical Wnt protein secretors maintaining FE patency

Having determined the importance of Wnt activity and Axin2+ cells at the FE, we suspected that Axin2+ cells may be an essential source of on-site Wnts. To assess this, *Wls*-depletion in Axin2+ cells was performed using *Axin2^CreERT2/+^Wls^f/f^* mice. At 1-week, histology confirmed an obstruction in both SG types, although lacking marked hyperplasia **(Fig. 4 A-D, S3 A-J)**. The obstruction was established without a marked immune cell response **(Fig. 4 E-H)**, with basal hyperproliferation from 9.5±1% in controls to 20.6±3% in experimental animals **(Fig. 4 I-L, Y).** K6A expression was elevated **(Fig. 4 M-P)**, signs of aberrancy, although slight, were more evident in the spinous and granular layers in the MG-FE rather than the SSG-FE **(Fig. 4 Q-T)**, cornification was lost within the FE in both SGs with signs of MGD in the MG-FE **(Fig. 4 U-X).** These findings indicate a similar obstruction in both SGs to those seen upon global-*Wls*-depletion, inferring Axin2+ cells are an essential source of Wnts ensuring FE patency. Dermal lengthening of the SSG associated hair had also occurred **(Fig. 4 A’-C’)**.

**Figure 4.**
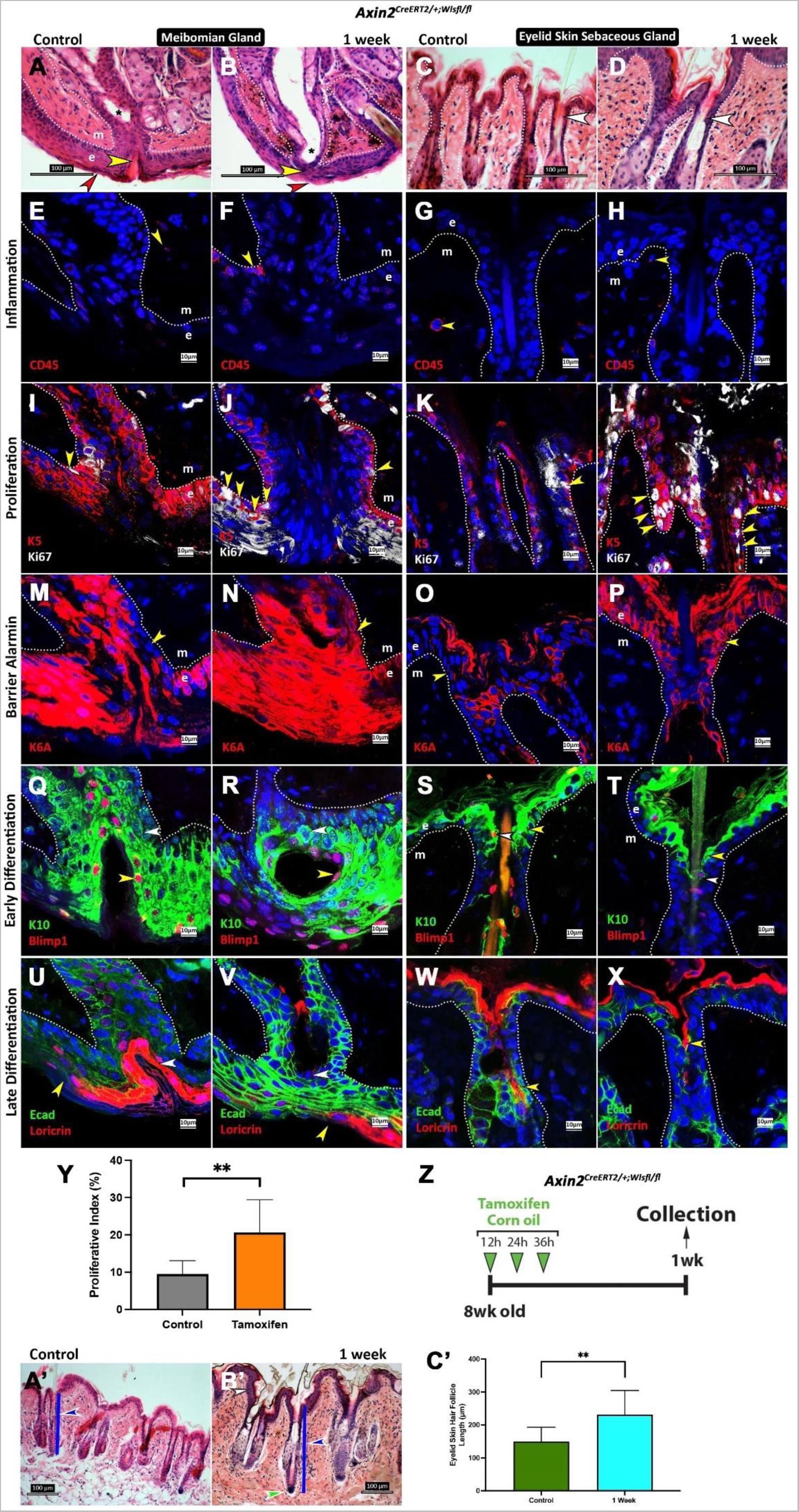
Depleting Wls in Axin2 stem/progenitors reveals them as critical Wnt protein secretors maintaining FE patency. **Z** Tamoxifen (experimental) and corn-oil (control) time course administration to 8-week-old *Axin2^CreERT2/+^Wls^f/f^*littermate mice and corn-oil controls (n=3 for 1-week timepoint). **A-D** Sagittal H&E. **A** Control MG-FE shows zone of keratinization posterior to MG-FE **(red)** and healthy cornification within the MG-FE **(yellow)**. **B** 1-week experimental shows a likely obstruction **(yellow)** lacking obvious hyperplasia but with anterior migration of the zone of keratinization **(red)**. **C** Control SSG-FE shows thin epithelia with space between it and the hair-shaft **(white)**, **D** experimental shows thickened epithelia with a loss of space. **E-X** Immunofluorescence. **E, G** Control samples show few to no CD45+ cells at SSG-FE **(yellow)**. **F, H** Experimental may show mild increase in CD45+ cells. **I, K** Control – few proliferating (Ki67+) basal (K5+) cells **(yellow)** at the SSG-FE. **J, L** Experimental – Increase in number of Ki67+ proliferating cells. **Y** Quantification of proliferation, all counts derived from n=3 animals at 1-week and subjected to unpaired student t-tests (error bars = SD). **M, O** Control K6A expression is sparsely present **(yellow)**. **N, P** Experimental samples showed an increase in K6A expression. **Q** Control MG-FE shows an orderly K10+Blimp1- and Blimp1+ expression rendering a patency which is **R** lost in the experimental sample. **S, T** Differences in the SSG-FE were less apparent. **U** Control Loricrin expression had a healthy posterior (yellow) and proximal (white) extension in regard to the MG-FE, **W** with an extension seen reaching the SG in the SSG-FE (yellow). **V-X** This was displaced distally in experimental samples **(yellow)**. **A’** Control shows hair-follicles of normal length in the dermal portion (blue). **B’** 1-week samples show elongation of the dermal portion of the hair-follicle, particularly at the bulb region (green). **C’** Quantification of the increase in length of the dermal portion of the hair-follicle. Scale bars and markers per image are indicated.

### 5. Constitutive Wnt activation in Axin2+ stem/progenitor cells results in FE keratin plugging and Axin2 clonal clustering at the PF

Having identified Axin2+ cells and their importance, we then assessed the effect of constitutive Wnt activation within Axin2+ cells using *Axin2^CreERT2/+^Ctnnb1^floxE3/+^;ROSA26^mTmG/+^*mice. 19-days was a terminal timepoint as mice were evidently poorly. Inflammation of the eyelids was evident **(Fig. 5 A, B)**. Histology revealed dense keratin plugging within the FE of both SG types, a proximally enlarged MG-FE and signs of neoplasia specifically at the PFs **(Fig. 5 C-F)**.

**Figure 5.**
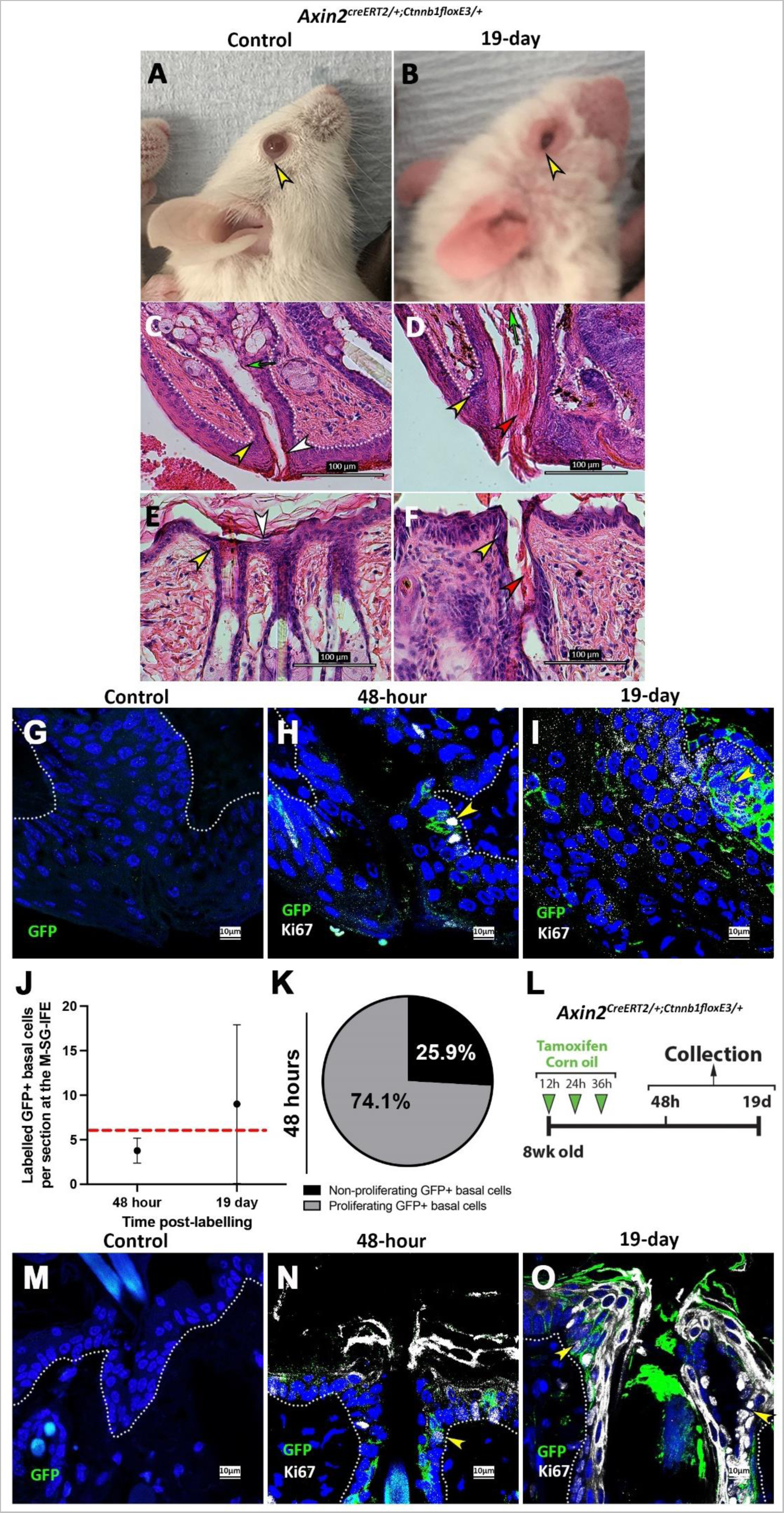
Constitutive Wnt activation in Axin2 stem/progenitor cells results in FE keratin plugging and Axin2 clonal clustering at the PF. L. Tamoxifen (experimental) and corn-oil (control) time course administration to 8-week-old *Axin2^CreERT2/+^Ctnnb1^floxE3/+^*littermate mice and corn-oil controls collected at a 48-hour and 19-day timepoint (n≥3 for both timepoints). **A** Control mice have normal healthy eyelids, fur, pale pink ears, and whiskers. **B** 19-day experimental mice are poorly with visible piloerection, fur, and whisker loss, along with eyelid snout and ear inflammation. **C-F** Sagittal H&E. Control shows healthy **C** MG-FE and **E** SSG-FE epithelium with an even and well-spaced distribution of basal and spinous nuclei **(yellow)** supporting a healthy granular and cornified layer **(white)**. Experimental shows abnormal **D** MG-FE and **F** SSG-FE epithelium with basal cell nuclei appearing dysplastic **(yellow)** and a keratin plug obstructing the SG lumen **(red)**. Immunofluorescent lineage tracing of *Axin2^CreERT2/+^Ctnnb1^floxE3/+^* G-I MG-FE and **M-O** SSG-FE. **G, M** Control indicates no line leakage. **H, N** 48-hour shows several basal Ki67+ cells some of which are GFP+ **(yellow)**. **I, O** 19-day shows large clusters of Ki67+ and GFP+ cells. **J** GFP+ basal cell counts at both time points in the MG-FE. K Number of GFP+Ki67+ vs GFP+Ki67-cells at 48-hours. All counts were derived from n=3 animals and subjected to unpaired student t-tests (error bars = SD). Scale bars per image are indicated.

Lineage tracing of the Axin2+ GFP+ clones in this line was then investigated. At 48-hours, markedly fewer GFP+ cells were detected than in homeostasis, however when they were detected, they were found in clusters and never solitary in both FEs and approximately 74.1±3% of these cells were proliferating. At 19 days, large proliferative clusters created greater variation in proliferative basal cell counts due to the lack of cell dispersion **(Fig. 5 G-O)**.

### 6. Constitutive Wnt activation in Axin2+ cells causes obstruction due to hyperkeratinisation fuelled by increased proliferation and differentiation

We then assessed the effect of constitutive activation of Wnt signalling in Axin2+ cells using molecular markers. It resulted in a milder localised increase in proliferation at 48-hours, with a marked generalised increase in proliferation at 19-days and a notable increase in the K5+ basal compartment size in the latter **(Fig. 6 A-D, N-P)**. Suprabasal expression of K6A showed a progressive increase **(Fig. 6 E-G, Q-S)**. Spinous and granular compartment sizes had approximately doubled in cell layer numbers by 19-day each indicating several more cells to be undergoing differentiation **(Fig. 6 H-J, T-V)**. At 48-hours in the MG-FE, cornification curiously showed a marked reduction with signs of MGD, while the SSG-FE showed a slight reduction, with sometimes no discernible change in the FE. At 19-days a dense keratin plug was present in the MG-FE indicating constitutive activation of Wnt signalling in Axin2+ cells is sufficient to increases cell proliferation and differentiation, leading in turn to hyperkeratinisation and blockage in a constrained space **(Fig. 6 K-M, W-Y)**.

**Figure 6.**
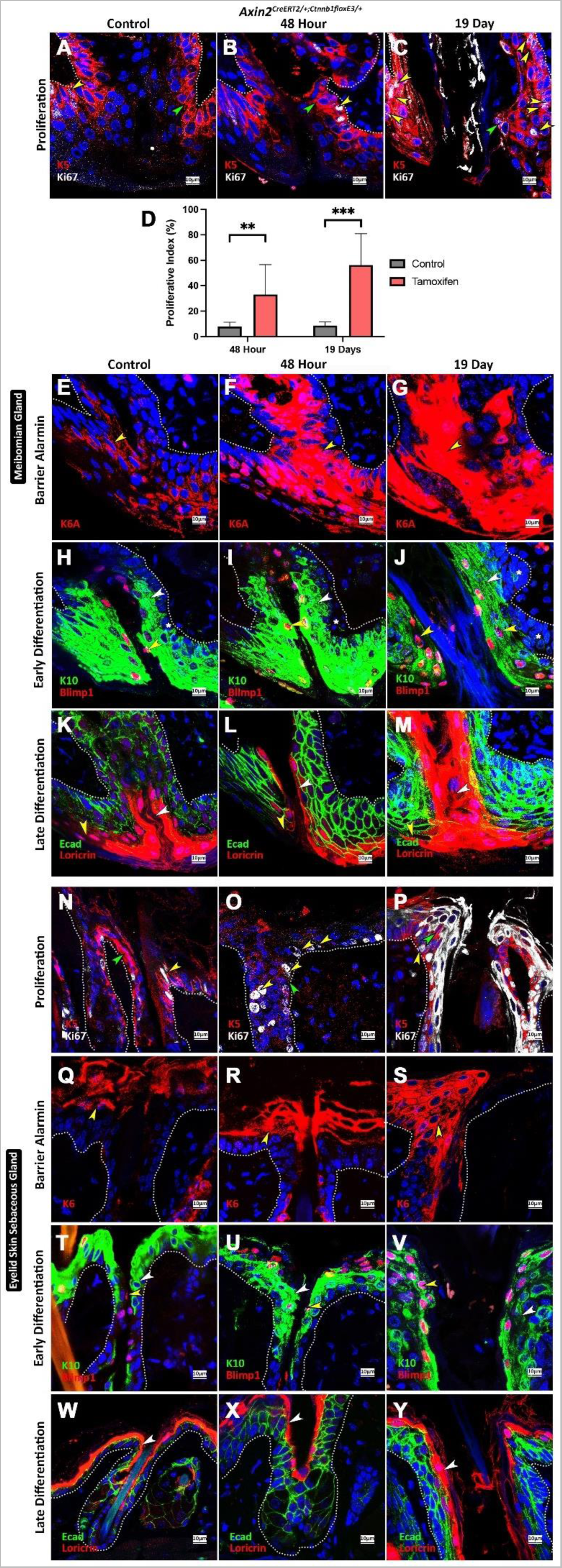
Constitutive Axin2 Wnt activation obstruction is due to hyperkeratinisation fuelled by increased proliferation and differentiation. 8-week-old *Axin2^CreERT2/+^Ctnnb1^floxE3/+^* littermate mice and corn-oil controls collected at a 48-hour and 19-day timepoint (n≥3 for both timepoints). Immunofluorescence at the **A-C, E-M** MG-FE and N**-Y** SSG-FE. **A, N** Control shows few Ki67+ proliferating cells (yellow) in a narrow K5 basal compartment (green). B, O 48-hour shows an increase in Ki67+ cells and possibly the size of the MG-FE K5 compartment. C, P 19-day shows increased proliferation and K5+ cell numbers. D Quantification of proliferation at the MG-FE. All counts were derived from n=3 animals per timepoint and subjected to unpaired student t-tests (error bars = SD). **E, Q** Control K6A (barrier alarmin) expression is sparse. **F, R** 48-hour and **G, S** 19-day K6A expression is elevated (yellow). **H, T** K10+Blimp1-expression labels a healthy spinous (white) and K10+Blimp1+ (yellow) in the control (asterisk in MG-FE indicates K10-basal layer at MG-FE-PF). **I, U** 48-hour and **J, V** 19-day samples show expansion in spinous and granular layers (underlying basal layer expansion in MG-FE-PF). **K, W** Control shows normal Loricrin expression within the FEs (white) and extending posteriorly in MG-FE (yellow). **L, X** 48-hour shows a mild reduction in Loricrin expression at the FEs, the MG-FE has anterior displacement of the zone of keratinization (yellow). **M, Y** 19-day shows a dramatic increase in Loricrin expression within the FE lumen, however, the MG-FE anterior displacement remains. Scale bars and markers per image are indicated.

### 7. Constitutive Wnt activation in Axin2+ cells tips cells in the PF toward a malignant neoplastic fate

Given the influential role of Wnt signalling in tumorigenesis, we then further investigated potential histological signs of neoplasia at the FE at 19-days in mice with constitutive Wnt activation in Axin2+ cells. Interestingly, neoplasia was noticeable in FEs including that of the eyelash at the PF. Notable features include well defined round to oval, elongated and sometimes irregular cell nests filled with cells carrying hyperchromatic nuclei and a scant cytoplasm. These and the often-seen retraction artefacts are hallmarks of basaloid neoplasia architecture **(Fig. 7 A-G)**.

**Figure 7:**
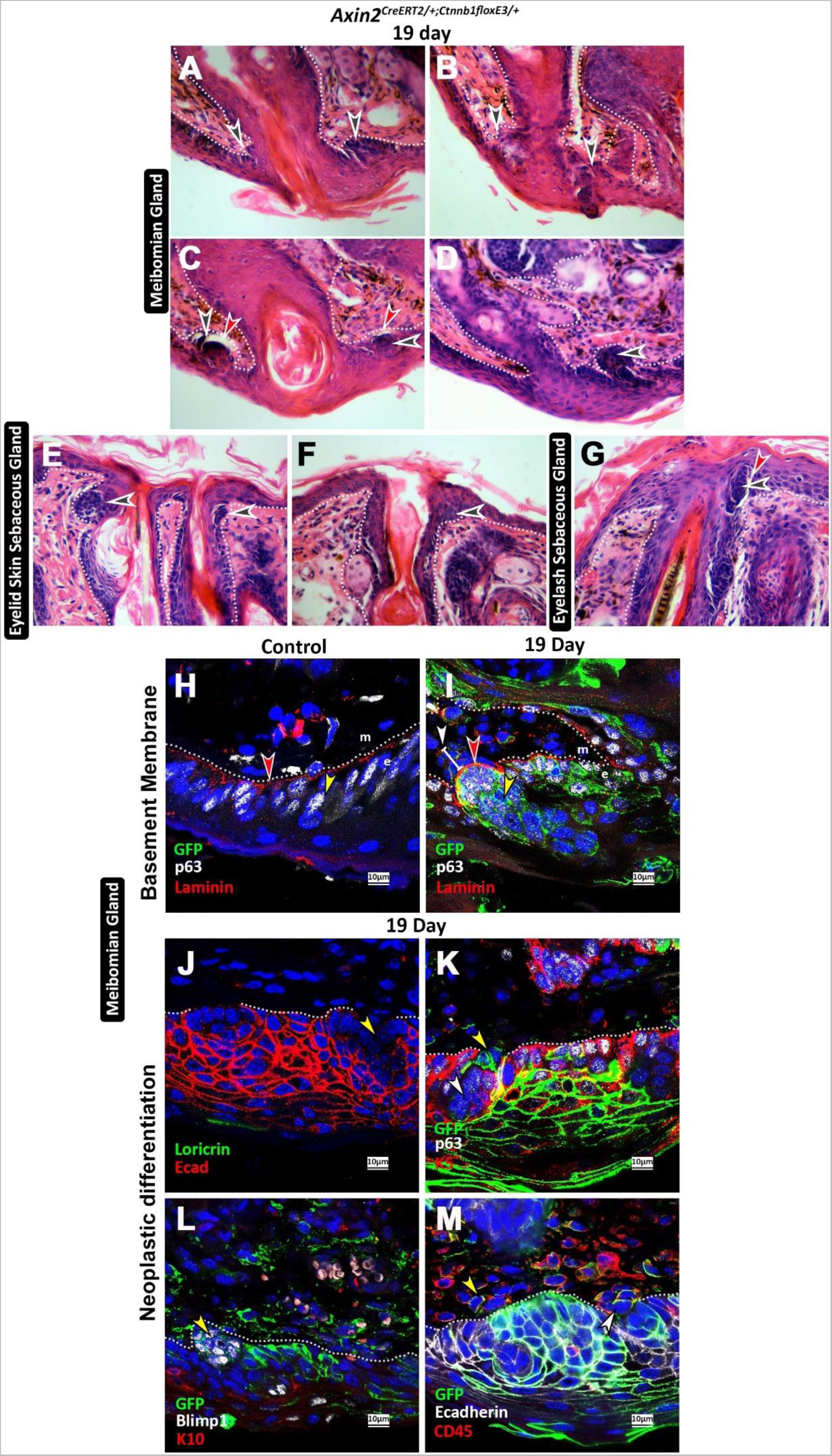
Constitutive Wnt activation appears to tip cells in the basal FE-PF toward a malignant neoplastic fate. 8-week-old *Axin2^CreERT2/+^Ctnnb1^floxE3/+^*littermate experimental mice collected at a 19-day timepoint (n>3). Sagittal H&E at the **A-D** MG-FE and **E-G** SSG-FE (including the eyelash SG) showing differing views of the suspected neoplastic clusters. Cell nests that are always filled with irregular spaced hyperchromatic nuclei with scant cytoplasm, hallmarks of basaloid architecture **(black)** with retraction artefacts **(red)**. **H-M** Immunofluorescence at the MG-FE in the region of the FE, section may be a sequential slice to the MG-duct. **H** *Axin2^CreERT2/+^Ctnnb1^floxE3/+^* littermate controls (n>3) show the basal cell nuclei labelled by p63 to be devoid of clustering **(yellow)**. Laminin **(red)** shows a gentle spotty appearance as it lines the basement membrane at the mesenchyme **(m)** – epithelium **(e)** boundary. **I** 19-day experimental samples show Axin2 p63+ basal clones **(yellow)** clustering. An aberrant increase in Laminin **(red)** seen at a part of the basement membrane that appears to irregularly invade the mesenchymal space. In the mesenchyme beyond the basement membrane, a mass of cells, some Axin2 clones, are seen extending proximally **(white line and arrow)**. **J-M** 19-day experimental samples. **J** A loss of the pan-epithelial cell marker E-cadherin in the centre of clusters **(yellow)**. **K** An Axin2 clone **(yellow)** devoid of p63 nuclear expression appearing to either break off from or interact with the adjacent cluster **(white)**. L Basal cell clusters aberrantly upregulate Blimp1 (yellow), clustered cells do not express the epithelial differentiation marker K10. M Several Axin2+ mesenchymal clones that are not but interact with CD45+ immune cells. Scale bars and markers per image are indicated.

Molecularly, Laminin, a marker of the basement membrane, showed increased expression suggesting malignancy of clusters at the interface with the mesenchyme **(Fig. 7 H, I)**. The tumorigenic clusters were also seen to lose epithelial characteristics and lacked differentiation characteristics (K10-, Loricrin-) **(Fig. 7 J)**. Intriguingly, Axin2+ clones were detected at the tumour-stromal front either breaking off or interacting with the tumorous lesion **(Fig. 7 K)**. Aberrant Blimp1 basal expression was also seen within the clusters **(Fig. 7 L)**.

## Materials and Methods

### Animal models

Transgenic mice (maintained on mixed background) and wild-type mice (CD-1 background) were housed in controlled conditions at the King’s College London Biological Services Unit. Wild-type CD1 mice were bred in-house. *pCag^creERT2/+^* (Hayashi & McMahon, 2002)*, Axin2^creERT2/+^* (R. van Amerongen *et al.*, 2012), *R26^mTmG/+^* ^57^, *Wntless^flox-flox^ ^(fl/fl)^* ^58^, *β*-catenin^lox(ex3)^ *(Ctnnb1^floxE3^*) (Harada *et al.*, 1999), *ROSA26-eGFP ^-DTA^* ^59^ and *TCF/Lef:H2B-GFP* ^54^ have been previously described. Mice were culled by UK Home Office approved exposure to carbon dioxide gas in a rising concentration (followed by femoral artery secondary confirmation) to ensure preservation of ocular structures by trained and approved individuals. Use of genetically modified mice was approved by the local GMO committee at King’s College London, under personal and project licences in accordance with the Animal (Scientific Procedures) Act of 1986, UK. Pig ocular samples were obtained from the Royal Veterinary College from 4-month-old juvenile healthy male pigs culled in accordance with home office guidelines. This project conforms with ARRIVE (Animal Research: Reporting of *In-Vivo* Experiments).

For all inducible mouse lines, intraperitoneal injections of tamoxifen (Sigma T5648) dissolved in corn oil at 20mg/ml with 10% ethanol were administered, at a dose of 0.15 mg/g body weight, 12-hourly three times and collected at the indicated time points after the first injection. For all experiments, both female and male mice were non-preferentially used.

### Immunofluorescence

8µm deparaffinized sections were rehydrated and heat-induced antigen retrieval was performed using 10mM Sodium Citrate (pH 6). Sections were then blocked in 1% BSA, 0.1% Triton-X, 1% goat serum and 0.5% of TSA kit (Perkin-Emler) blocking powder. Primary antibody incubation left at 4°C overnight. Sections were imaged with a Leica TCS SP5 or a Zeiss LSM800 Airyscan. To ensure true and specific protein expression, procedural controls were employed. Image processing and

### Immunofluorescence Antibodies

The following primary antibodies were used.

| **Name** | **Species** | **Company** | **Reference** | **Concentration** |
| --- | --- | --- | --- | --- |
| GFP | Chicken | Abcam | 13970 | 1/300 |
| E-cadherin | Mouse | Abcam | 76055 | 1/200 |
| E-cadherin | Rabbit | Abcam | 15148 | 1/200 |
| Endomucin | Rat | Abcam | 106100 | 1/100 |
| Ki67 | Rat | Invitrogen | 14-5698-82 | 1/150 |
| CD45 | Rabbit | Abcam | 10558 | 1/100 |
| K6A | Rabbit | BioLegend | 169P | 1/200 |
| K10 | Rabbit | Abcam | 9026 | 1/200 |
| K19 | Rabbit | Abcam | 52625 | 1/100 |
| K5 | Rabbit | Abcam | 52635 | 1/100 |
| P63 | Mouse | Abcam | 735 | 1/100 |
| Blimp1 IV | Rat | Invitrogen | 6D3 | 1/150 |
| Loricrin | Rabbit | Abcam | 85679 | 1/200 |
| Laminin | Rabbit | Sigma | L9393 | 1/30 |

The following secondary antibodies were used.

| Name | Species | Company | Reference | Conc. |
| --- | --- | --- | --- | --- |
| Alexa Fluor 633 | Goat anti-rat | ThermoFisher | A21094 | 1/200 |
| Alexa Fluor 633 | Goat anti-rabbit | ThermoFisher | A21073 | 1/200 |
| Alexa Fluor 568 | Goat anti-rat | ThermoFisher | A-11077 | 1/200 |
| Alexa Fluor 568 | Goat anti-rabbit | ThermoFisher | A-11011 | 1/200 |
| Alexa Fluor 488 | Goat anti-chicken | ThermoFisher | A11039 | 1/200 |
| DyLight 488 | Goat anti-rabbit | VECTOR | DI-1488 | 1/200 |
| DyLight 549 | Goat anti-rabbit | VECTOR | DI-1549 | 1/200 |
| DyLight 594 | Goat anti-rabbit | VECTOR | DI-1594 | 1/200 |
| DyLight 488 | Goat anti-mouse | VECTOR | DI-2488 | 1/200 |
| DyLight 549 | Goat anti-mouse | VECTOR | DI-2549 | 1/200 |
| DyLight 594 | Goat anti-mouse | VECTOR | DI-2594 | 1/200 |
| DyLight 649 | Goat anti-mouse | VECTOR | DI-2649 | 1/200 |

### Haematoxylin and Eosin staining histology

Eyelids were dissected and fixed overnight at 4°C in 4% paraformaldehyde (PFA), and then dehydrated through an ascending series of graded ethanol, before wax infiltration with paraffin wax at 60°C. Microtomy of 8 µm thick paraffin wax-embedded samples was performed prior to mounting on Superfrost® Plus slides. The slides were then stained with Haematoxylin and Eosin using standard techniques. Ehrlich’s Haematoxylin (Solmedia), Acid Alcohol (1%HCl in 95%IMS), 0.5%Eosin Y, 2mM Acetic Acid (Sigma-Aldrich). Staining was performed either by hand or using the Gemini AS Slide Stainer. Care was taken to store, monitor and maintain quality of all solutions over time. A bright field Carl Zeiss Axioskop 2 plus (1051-078) with a Nikon DS-Fi2 camera were used to take pictures.

### RNAscope

Axin2 expression RNAscope was done on 5μm thick formalin fixed paraffin embedded sections using the RNAscope 2.5 HD Reagent Kit-RED assay (Advanced Cell Diagnostics) following standard manufacturer’s instructions ^60^. Probe M. musculus Axin2 (Advanced Cell Diagnostics 400331) was used. A bright field Carl Zeiss Axioskop 2 plus (1051-078) with a Nikon DS-Fi2 camera were used to take pictures.

### Whole mount Oil-Red-O (ORO) staining

Following overnight PFA fixation, samples were transferred to labelled Eppendorf tubes and rinsed for 5-minutes 3-times with 1xPBS/0.5% Tween-20. Lipid staining was performed with a mixture of 300μl of 0.5% ORO in 100% isopropyl alcohol and 200μL of distilled water for 15 minutes. The samples were then rinsed twice with 1x PBS-Tween for 5-minutes each. Samples were then immersed into 60% isopropyl alcohol twice for 5-minutes each before a brief PBS-Tween rinse after which 4% PFA fixation was performed for 10 minutes. Samples were stored in 100% glycerol at 4°C. A Leica MZ FLIII was used to image.

### Proliferation index statistical analysis

All proliferating index counts were performed by counting the total number of basal cells in an image (y) and counting the total number of Ki67+ cells (x). Then x was divided by y and multiplied by one hundred to offer a percentile. A mean and standard deviation of each percentile was generated and graphed. The p-value was calculated using an unpaired two tailed t-test and significance then indicated in the graph.

### Cell counting

A minimum of three randomly selected images per mouse per experiment to a minimum of *n=3* mice were analysed. A magnification of x63 for confocal microscopy images, on non-overlapping sections, were used. In cases where additional images were introduced for counts, a balance was maintained between all *n* samples.

## Discussion

### The FE is a distinct epidermal niche

While the IFE has been widely studied, recent insightful work by Roy *et al.* (2016), who studied the dorsal skin of mice, showed that progenitor cells closer to a hair experienced increased proliferation compared to more distant cells. This led to the suggestion of a progenitor cycling model that is differentially regulated depending on distance from a hair. This finding has perhaps for the first time suggested that the IFE, belonging to the same skin type, behaves differently depending on distance to an appendage within it. Here, we formally characterised the FE of the MG and SSG and showed that the FE is structurally, functionally, and molecularly distinct to the IFE. We show that the FE has its own stem cell population/niche, and is markedly affected by Wnt signalling aberrancy, resulting in contrasting forms of epidermal obstructions at the FE that likely affect SGs secretions and skin/cornea barrier protection.

We first identified that a long-lived Axin2 stem/progenitor cell population is responsible for Wnt protein secretion that ensures FE renewal through regulated proliferation and differentiation thus maintaining follicular patency. These cells reside in the PF pouch-like region observed in murine and porcine histology. The PF has rare *TCF/Lef:H2B-GFP+* cells, and is the site of neoplastic malignancy, from which we can infers that the site is a niche.

### Wnt signalling loss leads to FE dysregulation and clinical obstructive disease

Upon *Wls*-depletion, we found a unique obstruction at the FE prior to inflammation with marked hyperproliferation, hyperplasia, and inhibited terminal differentiation accompanied by eyelash loss. Current follicular obstructive pathologies do not account for such a mechanism in disease progression although several clues to their existence exist. Augustin and colleagues (2013) showed that murine *Wls*-depletion resulted in phenotypes resembling psoriasiform dermatitis in the IFE and confirmed down-regulation of *Wls* in psoriatic patient biopsies. Psoriasis manifests as itchy, scaly patches of epidermis undergoing hyperproliferation and swelling. Past work has shown psoriasis to have epithelial hyperplasia underpinned by pathological hyperproliferation ^50,52,61–66^. Additional signs include an increase in expression of early differentiation (spinous layer) markers ^67^, an inhibition of terminal differentiation at the late spinous layer ^51,68^, and abolishing of late differentiation markers ^67^. These signs are strikingly similar to our findings upon global *Wls*-depletion inferring SG obstructions via inhibited differentiation in patients with psoriasis caused by *Wls* downregulation. Furthermore, Iizuka *et al.* (2004) likened this pathogenic IFE phenomenon to a wound healing response ^69,70^, with psoriatic lesion improvement occurring after an extended turnover time as late differentiation markers attempt to catch up ^67^. While this catch-up opportunity for healing may occur at the open-plan IFE, the contained FE may be less forgiving resulting in an irreversible obstruction.

Sebum inhibition is a characteristic of psoriasis and SG atrophy has been suggested as the cause although a mechanism is lacking ^71,72^. SG atrophy is theorised to arise via either genetic lipid synthesis inhibition ^71^ or due to a lack of sebocyte differentiation ^72^. Our data suggests the possibility of an initial obstruction due to abnormal epidermal function at the FE resulting in gland atrophy, a mechanism seen upon salivary gland ligation ^73^ or salivary stone formation in humans ^74^. Furthermore, it is worth remarking that although inflammation is the prime suspect in psoriatic gland atrophy, an epidermal cytokine-release site has remained elusive ^71,75^. Our study suggests that obstructions can precede an inflammatory response, and thus the pathogenic FE is a potential site for cytokine-release. The idea of the epidermis acting as a psoriatic inflammatory trigger is an ongoing debate in the field ^76^. Notably, in further support of SG obstructions being prevalent in psoriasis, in the eye, psoriatic patients are known to have MG obstructions ^77,78^ and thus decreased meibum flow ^79^. Our findings offer a novel likely mechanism here and warrant a review to determine if SSGs of psoriasis patients are also obstructed.

Intriguingly, in the larger scope of disease, psoriasis has been suggested as a pre-diabetic condition owing to its links to type II diabetes mellitus ^80–82^, where most type II diabetics are also known to suffer from MGD ^83^. Wnt signalling and diabetes mellitus type II also share links ^84–90^ thus warranting both the molecular (Wnt dependent) and cellular (no involvement of FE hyperkeratinisation) mechanisms underlying MGD in our experiments. SSG-FE obstructions have not been described in type II diabetes and may have been overlooked. Current MG obstruction management involves antibiotics, steroids, anti-inflammatories, hormonal therapy, meibum expression, ductal probing, heating devices, pulsed light, and neurostimulation ^91,92^. In light of our findings, the efficacy of these management techniques is likely lacking as they do not directly address hyperplasia and/or inhibited differentiation. Our findings warrant a clinical review of concerned diseases to rule out obstructions of this nature.

Hair loss can be a debilitating condition and is known to occur, with an often-unknown aetiology and pathogenesis in several skin conditions such as alopecia areata, psoriatic alopecia, androgenetic alopecia, among several others ^93^. FE obstructions are not recognised and/or ruled out as a contributing factor to any of the conditions causing hair loss. Psoriatic alopecia is clinically well-established with loss of visible hairs on the skin surface and subcuticular elongation, however, the mechanism at play is unknown ^71,94,95^. We observed a generalised loss of eyelashes and loss of a subset of eyelid hairs in mice upon *Wls*-depletion, suggesting the possibility of anomalous mechanics contributing to hair loss-defects. It is likely that terminal differentiation inhibition, as seen in our experiments, may implicate exposed and opposed spinous layer cells to adhere to each other and/or the hair shaft during obstruction, resulting in downward trapped growth with surface hair loss.

Constitutive Wnt/β-catenin activation in Axin2 stem/progenitor cells, resulted in a different form of obstruction to *Wls*-depletion at the FE. Hyperproliferation, increased cell input and thus differentiating cells resulted in keratin plugging of the SG-FE. Hyperkeratinisation and plugging is present in acne in the skin and MGD in the eyelid however lacking a substantial direct link to Wnt signalling. Lithium, a commonly used psychiatric drug and an established Wnt/β-catenin signalling activator ^96^ offers a useful clinical indicator that validates our findings. Among the several aggressive clinical side-effects of lithium, among the most common are acne ^97,98^, and dry eye disease via tear-film dysfunctions ^99,100^. Our findings suggest the latter feature to be due to obstructive MGD and warrants a clinical review of this. A claimed gold standard of acne therapy is oral isotretinoin treatment, which has an anti-keratinising action at the cornified layer and aggressively limits sebum production by inducing SG atrophy (Allenby et al., 1993; Dalziel et al., 1987; Dréno, 2017; Fogh et al., 1993; Geiger et al., 1996; Layton, 2009; Ott et al., 1996; Plewig et al., 2004; Tsukada et al., 2000; Vallerand et al., 2018). Isotretinoin has well-documented side effects including xeroderma, teratogenicity, psychiatric, gastrointestinal, and neurological symptoms ^111^. Markedly cheilitis affects 90% of patients on isotretinoin with dry eye also being a frequent symptom ^112^. Notably, retinoic acid (treats acne), is known to suppress Wnt/β-catenin signalling ^113^, and additionally, Wnt/β-catenin signalling activation is required for sebocyte differentiation ^114^, suggesting Wnt signalling suppression may be central to isotretinoin action and acne treatment as our data suggests in the mouse. These findings may aid drug development for acne treatment with fewer side effects.

### Activation of Wnt signalling and basaloid neoplasia at the FE

Aside from obstructions, a constitutive activation of Wnt signalling resulted in tumorigenesis owing to Axin2+ cell hyperproliferation appearing to cause hyperproliferation in neighbouring GFP-cells specifically at the FE-PF, a further niche-like characteristic. Given that day 19 was a terminal time point, advantageous late-stage diagnostic features of the cancer were not possible although histology revealed the malignant neoplasia to have retraction artefacts and these are a reported feature of extracellular matrix degradation ^115^. Furthermore, molecularly, we found the basement membrane at the site to have laminin overexpression which is linked to uncontrolled cell proliferation, invasion, and metastasis ^116–118^. Basal neoplastic cells appeared to lose epidermal characteristics and exhibited ectopic Blimp1 expression. Blimp1/PRDM1 is an immune system transcriptional repressor ^119^ and regulates epidermal keratinocyte terminal differentiation ^120^. In one of the most metastatic cancers, the pancreatic ductal adenocarcinoma (PDAC), hypoxia in the tumour microenvironment has been shown to promote Blimp1 expression and Blimp1 expression in PDAC acts as a switch that promotes metastasis while inhibiting cell division ^121^. Furthermore, Blimp1 is claimed to mediate epithelial-to-mesenchymal transition (EMT) through stimulation of a migratory phenotype and promotion of a tumour cell immune evasion respectively in breast cancer cells ^122^, lung cancer cell lines ^123^ and hepatocellular carcinoma ^124^. More recently Blimp1 has been identified as a marker predicting progression to a higher-grade disease for cervical intraepithelial neoplasia ^125^. Altogether our findings suggest the presence of an invasive form of malignant neoplasia. This warrants investigating Wnt/β-catenin activity and Blimp1 expression in eyelid neoplasia as it may have a diagnostic and/or prognostic value.

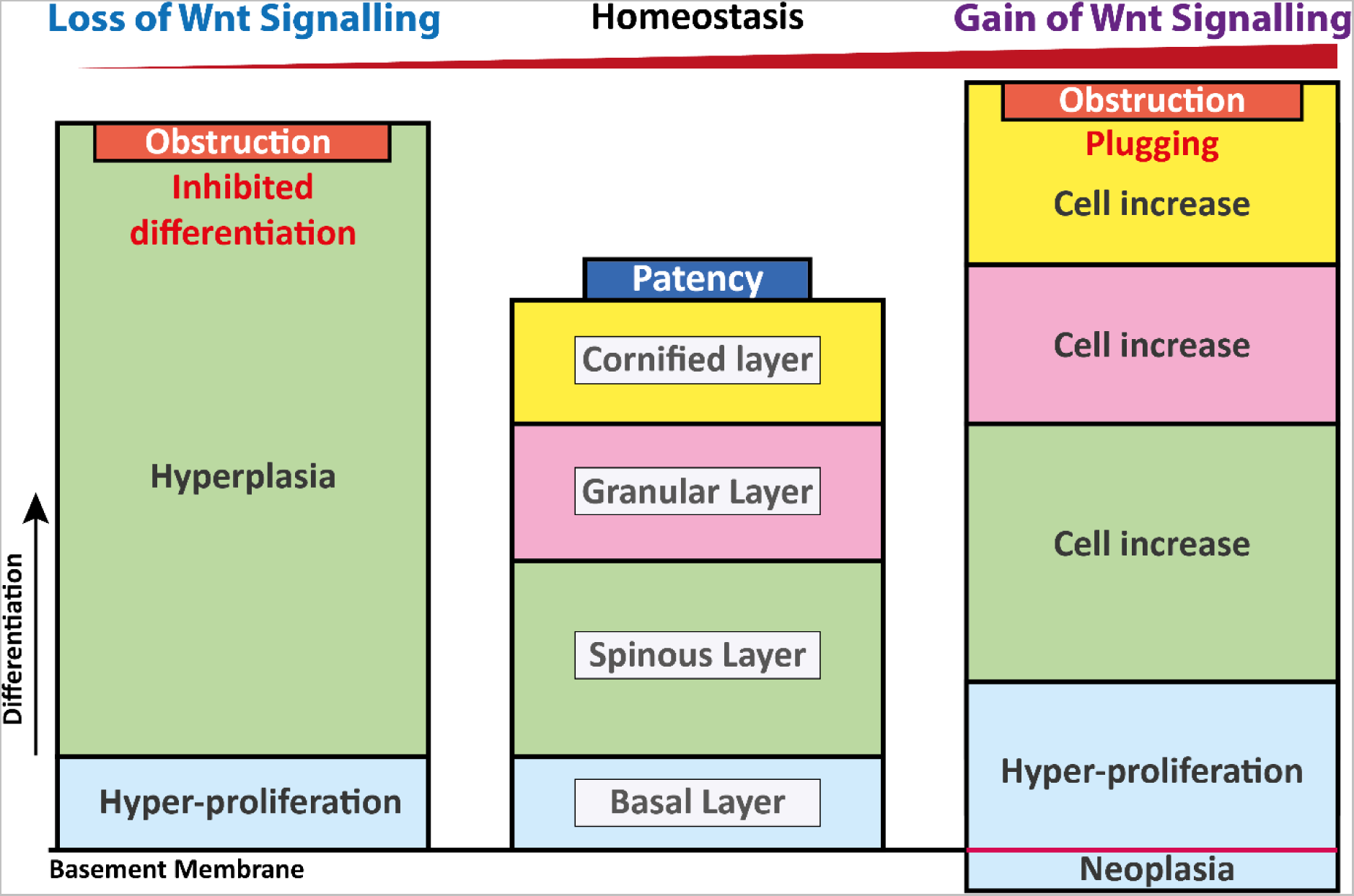

## Conclusion

Altogether, our findings place the FE as a pivotal site at the confluence of the IFE and the SG-follicle entrance and may also exist at the terminal end of other non-sebaceous dermal appendages such as the sweat glands. Adequate levels of Wnt signalling ensure FE homeostasis and thus prevention of obstruction onset in a multitude of diseases. We have identified two contrasting forms of obstructions, one via inhibited terminal epidermal differentiation (lack of keratinisation) and the other by hyperkeratinisation. While the latter is generally accepted, the former has currently not been documented to occur suggesting current therapies may fail due to misdiagnosis. Both mechanisms of obstruction have antero-displacement of the MCJ at the M-FE supporting current clinical diagnostic use of vital dye staining for the Line of Marx in MGD. Based on the known basic functions of keratin, such as smoothness and rigidity, these features may be lost rendering a lumen to have irregular surface characteristics thus inhibiting secretion flow. Furthermore, luminally opposing spinous layers may make contact, especially in hyperplasia, and may be more prone to adhering in the absence of cornification.

Overall, the FE is a key bottle neck region and a likely tipping point for several diseases that may be currently overlooked. Our findings warrant a review of epidermal conditions to rule out obstructions as inhibition of sebaceous secretions likely confound disease progression. Furthermore, the Wnt signalling pathway may offer novel more targeted therapeutic avenues to challenge FE disease pathophysiology in ophthalmology, dermatology, and oncology.

## Acknowledgements

We thank Prof Cynthia Andoniadou, Prof Paul Sharpe, and Prof Abigail Tucker for invaluable technical support. We thank Dr Emily Lodge and Dr Thea Willis for help with RNAscope. We thank Dr Nicholas Merrild for pig samples and Dr Jing Zhao for mouse samples. We thank Mr Alasdair Edgar and Ms Wai Lin Tsang for support with histology. S.S. was full funded by the King’s College London International Studentship.

## Author contributions

Conceptualisation, S.S. and I.M.; Methodology, S.S., I.M. and AA.; Software, S.S.; Validation, S.S. and A.A.; Formal Analysis, S.S.; Investigation, S.S.; Resources, I.M., G.P., A.A.,; Data Curation, S.S.; Writing Original Draft, S.S.; Writing Review & Editing, S.S., I.M., G.P., A.A.; Visualisation, S.S.; Supervision, I.M. and G.P.; Project Administration, S.S., I.M., and G.P.; Funding Acquisition, I.M. and G.P.

