## Supplementary Information for "Gate keeping the sebaceous gland"

Title

Supplementary figures

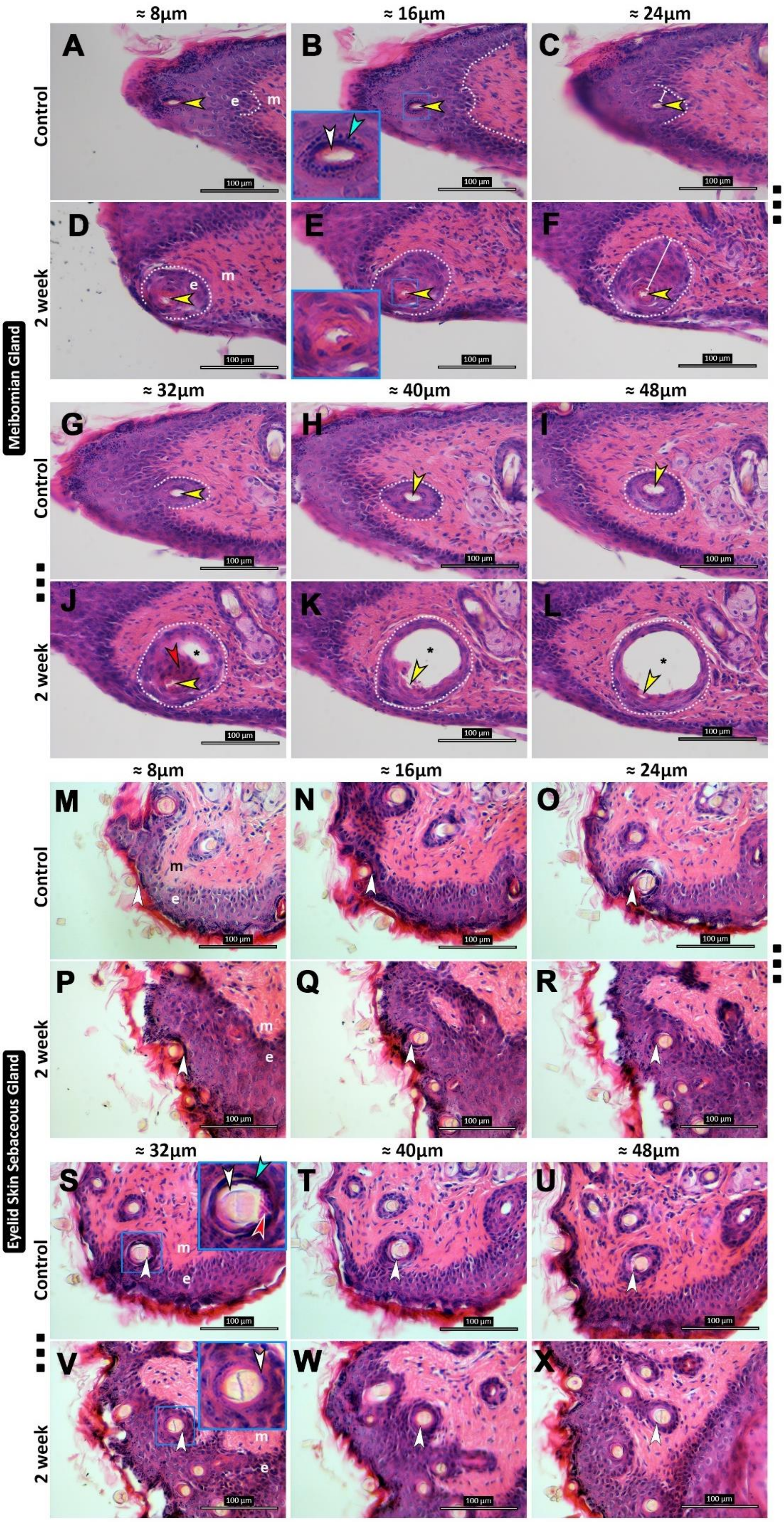

≈ 8µm
≈ 16µm
≈ 24µm
≈ 32µm
≈ 40µm
≈ 48µm
Control
2 week
Meibomian Gland
Eyelid Skin Sebaceous Gland
A
B
C
D
E
F
G
H
I
J
K
L
M
N
O
P
Q
R
S
T
U
V
W
X
e
m
100 µm

**Supplementary 1. Transverse H&E shows an obstructions associated with epithelial hyperplasia at the SSG-FE of both SGs upon global *Wls*-depletion.** H&E staining of transverse 8µm sequential sections at the MG-FE **(A-L)** and the SSG-FE **(M-X)** of *pCag*$^{CreERT2/+}$*Wls*$^{f/f}$ 2-week experimental and littermate control samples. Epithelium-**e**, mesenchyme-**m,** dotted line–boundary. **A-C, G-I** Control shows narrow but patent MG-FE lumen that is slightly divergent apically **(yellow arrow)** with healthy surrounding epidermal cell characteristics. **B (Blue rectangle** high magnification inset**)** Control 16µm height healthy granular **(cyan arrow)** and cornified layer **(white arrow)** seen. **D-F, J-L** 2-week shows a MG-FE obstruction **(red arrow)** with an abnormal lumen **(yellow arrow). E** Experimental has loss of granular and cornified layers. **F** Marked thickening of the MG-FE **(white line). J** Hyperplastic mass of cells **(red arrow)** interrupting the SSG-FE lumen and MG-duct **(asterisk). K** Marked MG-ductal dilation. **M-O, S-U** Control shows SSG-FE and its associated centrally placed hair-shaft with a discernible white ring of space between the two structures **(white arrow)**. **S** At 32µm**,** clear spacing present with healthy granular **(cyan arrow)** and cornified layer **(red arrow). P-R, V-X** 2-week shows a lack of space between structures **(white arrow)**. **V** 32µm, lack of space with altered granular and cornified layer**.** Scale bars per image are indicated.

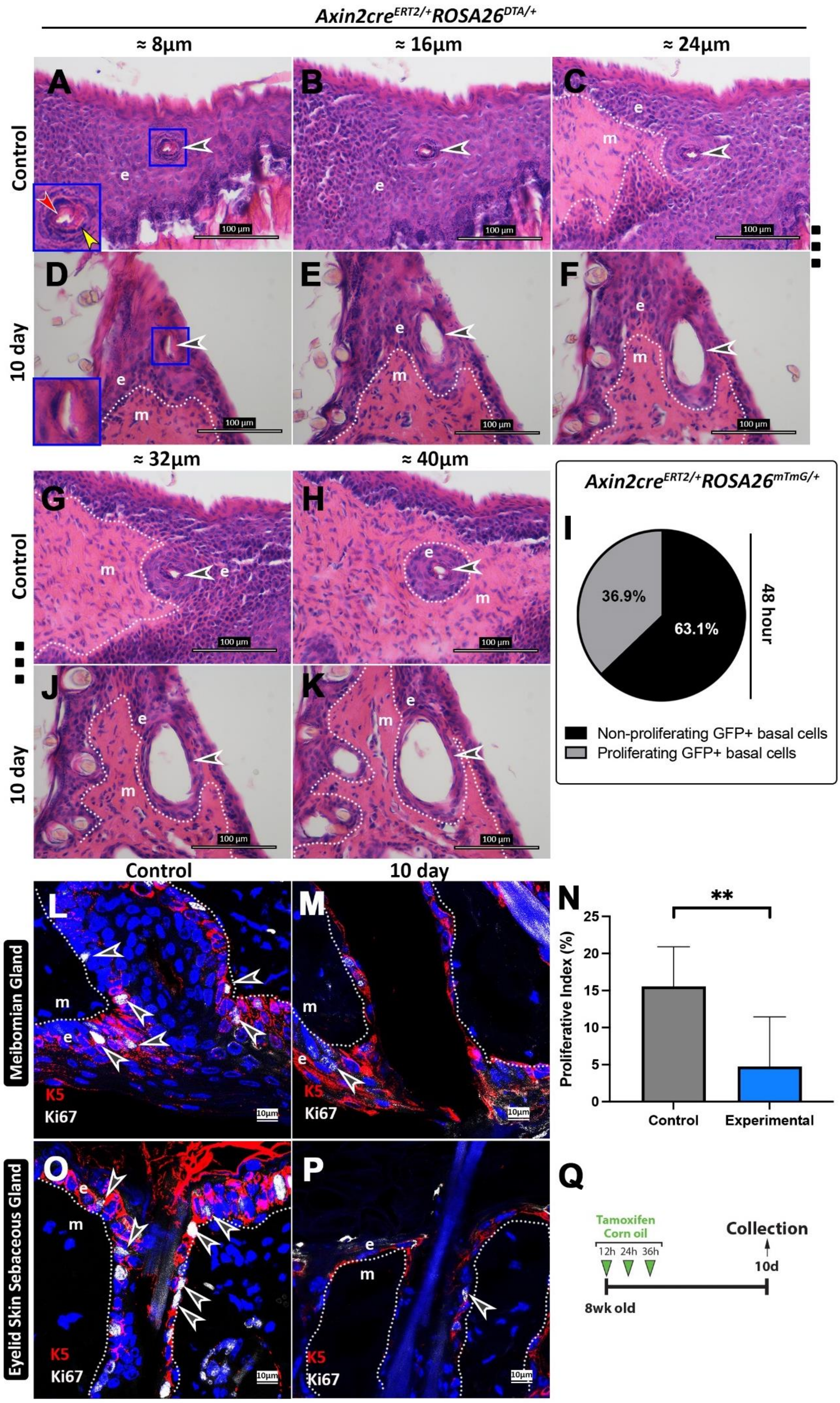

Axin2cre^ERT2/+ROSA26^DTA/+
≈ 8μm
≈ 16μm
≈ 24μm
A
B
C
Control
10 day
D
E
F
e
m
100 μm
≈ 32μm
≈ 40μm
G
H
J
K
Axin2cre^ERT2/+ROSA26^mTmG/+
I
36.9%
63.1%
48 hour
Non-proliferating GFP+ basal cells
Proliferating GFP+ basal cells
Control
10 day
L
M
N
**
Proliferative Index (%)
25
20
15
10
5
0
Control
Experimental
Meibomian Gland
Eyelid Skin Sebaceous Gland
K5
Ki67
10μm
O
P
Q
Tamoxifen
Corn oil
12h 24h 36h
Collection
10d
8wk old

**Supplementary 2. Axin2 ablation results in a loss of cell numbers in the SSG-FE with a marked reduction in basal proliferation. A-F, G, H, J, K** H&E staining transverse 8µm sequential sections from *Axin2*$^{CreERT2/+}$*ROSA26*$^{DTA/+}$ mice at the MG-FE. Luminal proximal divergence seen in experimental samples with thinning of the MG-FE wall (black). L, M, O, P Immunofluorescence showing K5 labelling basal cells and ki67 labelling proliferating cells. **M, P** 10-day show a marked reduction in K5+ and Ki67+ cell numbers at the SSG-FE. **N** Quantification of MG-FE proliferation. All counts derived from n=3 animals at 10-days and subjected to unpaired student t-tests (error bars = SD). **Q** Tamoxifen (experimental) and corn-oil (control) time course administration to 8-week-old *Axin2*$^{CreERT2/+}$*ROSA26*$^{DTA/+}$ littermate mice (n=3 for the timepoint). **I** Shows the percentage of GFP+ cells proliferating at 1-week in Axin2$^{creERT2+}$;ROSA26$^{mTmG/+}$ at the MG-FE (Refer to Fig. 3I).

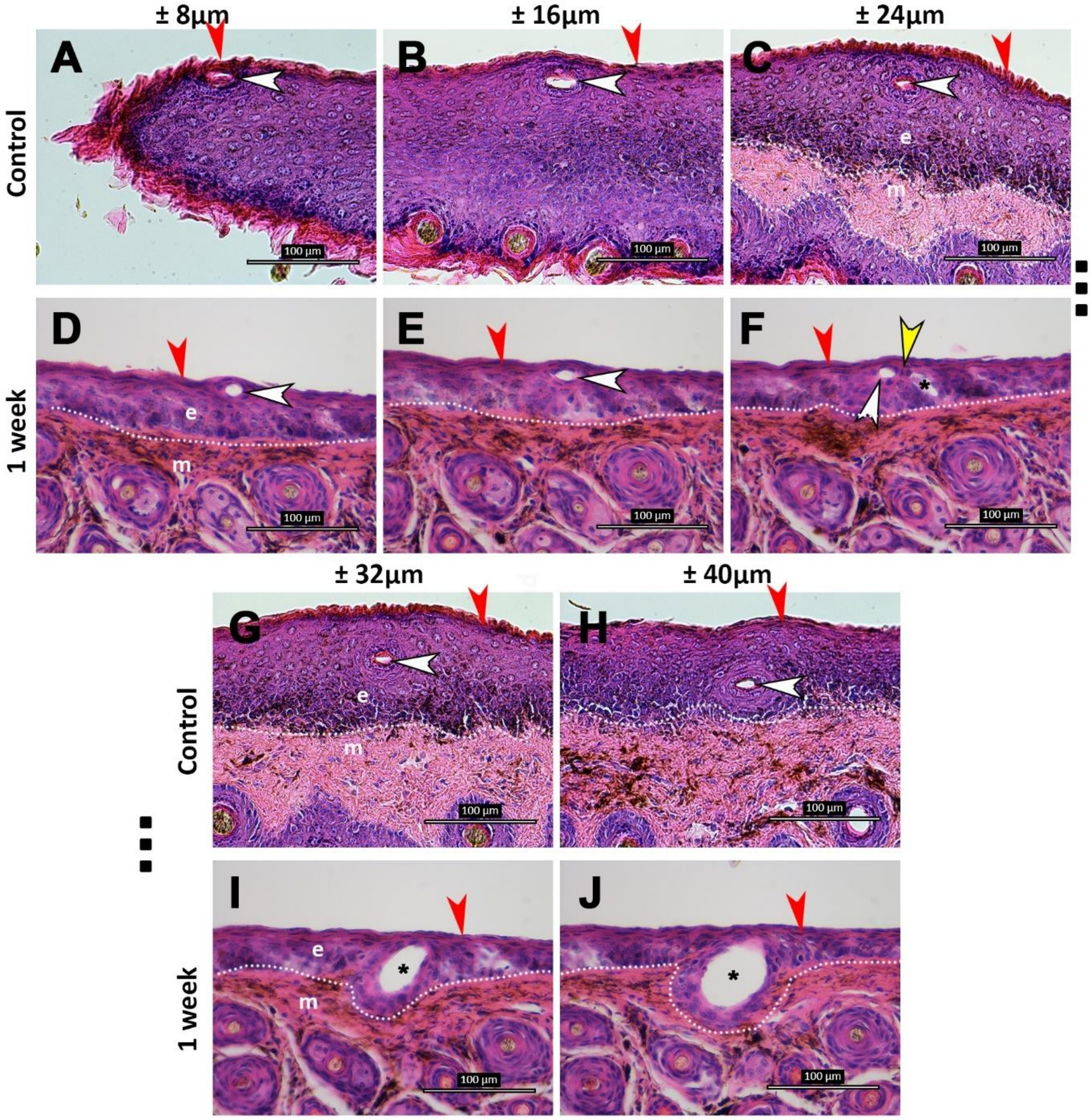


**Supplementary 3. Axin2+ *Wls*-depletion results in an obstruction likely linked to inhibited differentiation. A-J** H&E staining transverse 8µm sequential sections from 8-week-old *Axin2*$^{CreERT2/+}$*Wls*$^{f/f}$ littermate mice and corn-oil controls (n=3) at the MG-FE. **A-C, G, H** Controls show an ordered epidermis with a evident granular 'gritty' layer (white), cornification (red) and well-spaced evenly distributed nuclei in the epithelium. **D-F I, J** Experimental samples show a lacking granular and cornified layer, nuclei irregularly spaced (varying density) in the epithelium, and an obstruction present in **F**. Central duct (asterisk) is separated from the MG-FE (white) before dilating proximally. Scale bars and markers per image are indicated.
